# Genetic diversity within and between polyploid sugarcane (*Saccharum* spp.) families obtained via caryopsis using microsatellite markers and multicategory model

**DOI:** 10.64898/2026.08.28.747747

**Authors:** Luiz Gustavo da Mata Borsuk, Hugo Zeni Neto, Vitor Bialetzki Cristiano, Maria de Fátima Pires da Silva Machado, Claudete Aparecida Mangolin, Leticia Martins Montini, Joseli Cristina da Silva, Renato Frederico dos Santos

## Abstract

Genetic diversity analyses are essential for sugarcane (*Saccharum* spp.) breeding programs. Crossbreeding, based on genetic distances between parental plants, is a tool used to increase genetic variability and enhance plant selection; however, quantifying variation in highly polyploid species remains a challenge. The present study aimed to evaluate the diversity within and between 12 families of sugarcane derived from caryopses, analyzing 120 individual seedlings arranged in an augmented block design. Genotyping was performed using primers for 16 microsatellite loci, five simple sequence repeat (SSR) loci, and 11 expressed sequence tag-SSR (EST-SSR) loci. To accurately account for polyploidy, similarity calculations were performed using Bruvo’s distances among individuals and RST distances among the families. Analysis of molecular variance (AMOVA) indicated that most of the genetic variability was within families (72%), with only 28% found between them. This high level of intra-family variation demonstrates that a significant reservoir of genetic diversity remains available within the crosses. The highest genetic similarity was observed between the families RB986952 × RB986960 and RB036122 × RB03611, whereas the lowest genetic similarity was observed between the families RB97319 × RB966928 and RB106802 × RB855036. Although the evaluated families shared high genetic similarity, the pronounced genetic variation within them demonstrates a robust recombination potential, indicating that the genetic basis of sugarcane can be better explored using the high variability that already exists in the selection of desirable morpho-agronomic characteristics within the families. Furthermore, this study highlights the importance of using appropriate distances for diversity studies with codominant markers, such as microsatellites, in polyploid species.

## Author Summary

Sugarcane is a vital global crop for sugar and bioenergy production; however, creating new improved varieties is difficult because of its complex genetics. To make breeding programs more efficient and less costly, breeders need to understand where the genetic diversity lies, whether it is between different families of plants or hidden among siblings within the same family.

In this study, we investigated the genetic structure of 12 sugarcane families using molecular DNA markers and compared them with their agronomic traits in the field. We discovered that 72% of the total genetic variation was maintained within the families themselves, rather than between different families. Despite this general similarity across groups, we identified specific families that diverged significantly in terms of genetic architecture and agronomic traits.

These findings have practical implications for agricultural sciences. They demonstrated to sugarcane breeders that the true potential for crop improvement lies among siblings. By focusing on this within-family variation during the early stages of breeding, researchers can more effectively select plants with superior traits, ultimately accelerating the development of more productive and resilient sugarcane varieties.

## Introduction

Sugarcane (*Saccharum* spp.) is a major global commodity, occupying an area of over 26 million hectares in more than 130 countries, with a total production of over 1.5 billion tons. Brazil stands out in this scenario, accounting for approximately one-third of the world’s production, with a harvest of approximately 553 million tons [1]. Sugarcane plays a fundamental role in ethanol production. Brazil is one of the leading countries in renewable energy production, generating approximately 31.3 billion liters of ethanol, with 27.37 billion liters derived directly from sugarcane cultivation [2].

Despite its economic importance, genetic gains in breeding programs are consistently hindered by the species’ genomic complexity. Modern sugarcane cultivars used in commercial fields are highly polyploid and are derived from interspecific hybrids between *S*. *officinarum* (2n = 80) and its wild relative, *S. spontaneum* (2n = 40–128). The complexity of the sugarcane genome is due to its high number of chromosomes, ranging from 100 to 130, a genome size of approximately 10 Gb, and the occurrence of aneuploidy with a variable number of chromosomes in each homologous group [3–5].

According to [6], the study of genetic diversity is important for understanding the evolution of different populations at the ploidy level that exists among them, planning and implementing new crosses, and better utilization of the genetic base of sugarcane to create new cultivars for breeding programs. However, most of these studies characterize established commercial varieties or elite clones of advanced age [7–8], which are vegetatively propagated and have accumulated decades of physiological and somatic histories. Investigating genetic variability from the first day of the cycle using caryopses derived from controlled crosses remains largely unexplored. Evaluating genetic diversity at this initial stage is crucial for estimating the true baseline recombination potential of a population before any clonal selection or long-term vegetative propagation occurs.

To address these breeding challenges, molecular markers have been widely deployed as solutions for diversity characterization. Molecular markers, such as simple sequence repeat (SSR) loci, formed by simple repeated DNA sequences (also known as microsatellite loci), and expressed sequence tag-SSR (EST-SSR) loci, which correspond to SSR loci contained within expressed DNA sequences, have been successfully used to study genetic diversity in several sugarcane varieties [9–12].

Among these options, microsatellite markers have several advantages over other molecular markers because they are highly polymorphic and their inheritance is codominant, which enables discrimination between homozygous and heterozygous plants. They are multiallelic, amplified using PCR, and therefore require small amounts of DNA for analysis, and are highly reproducible [13,14]. However, in sugarcane, a highly polyploid species with different ploidy levels, SSR markers are treated as dominant and analyzed as binary, which may lead to a loss of information in the analysis.

To overcome this limitation, Bruvo’s genetic distance [15] can be implemented with multicategorical modeling to treat microsatellite markers as genuinely codominant markers in polyploid complexes. This approach enables the precise detection of multi-allelic heterozygosity and captures the exact genetic information per locus, avoiding the loss of information typical of binary data. Furthermore, in commercial fields, sugarcane is vegetatively propagated, which means that farms plant elite clones. Consequently, identifying superior and highly divergent parental genotypes for initial crosses is crucial, as any resulting genetic gain can be immediately fixed and multiplied by cloning.

Thus, the objective of the present study was to evaluate the genetic variability within and between 12 sugarcane families derived from caryopses generated in a breeding program. Using SSR and EST-SSR markers combined with Bruvo’s genetic distance, this study aimed to identify the most divergent genotypes and assist in the selection of crosses and subsequent stages of sugarcane genetic breeding programs.

## Results

The annealing temperatures of the primer pairs are listed in Table 1. Reactions were conducted to modify the primer temperatures because they did not align with those documented in the literature. The Touchdown program (TD-PCR) proved to be the most efficient for DNA amplification; 15 out of 16 primer pairs successfully amplified DNA from 12 families. A specific temperature of 53 °C was used only for primer SEGMS-1069. Amplification reactions using the *TD-PCR program* have been shown to be highly efficient for annealing SSR primers in sugarcane [11–12,16–18].

**Table 1.**
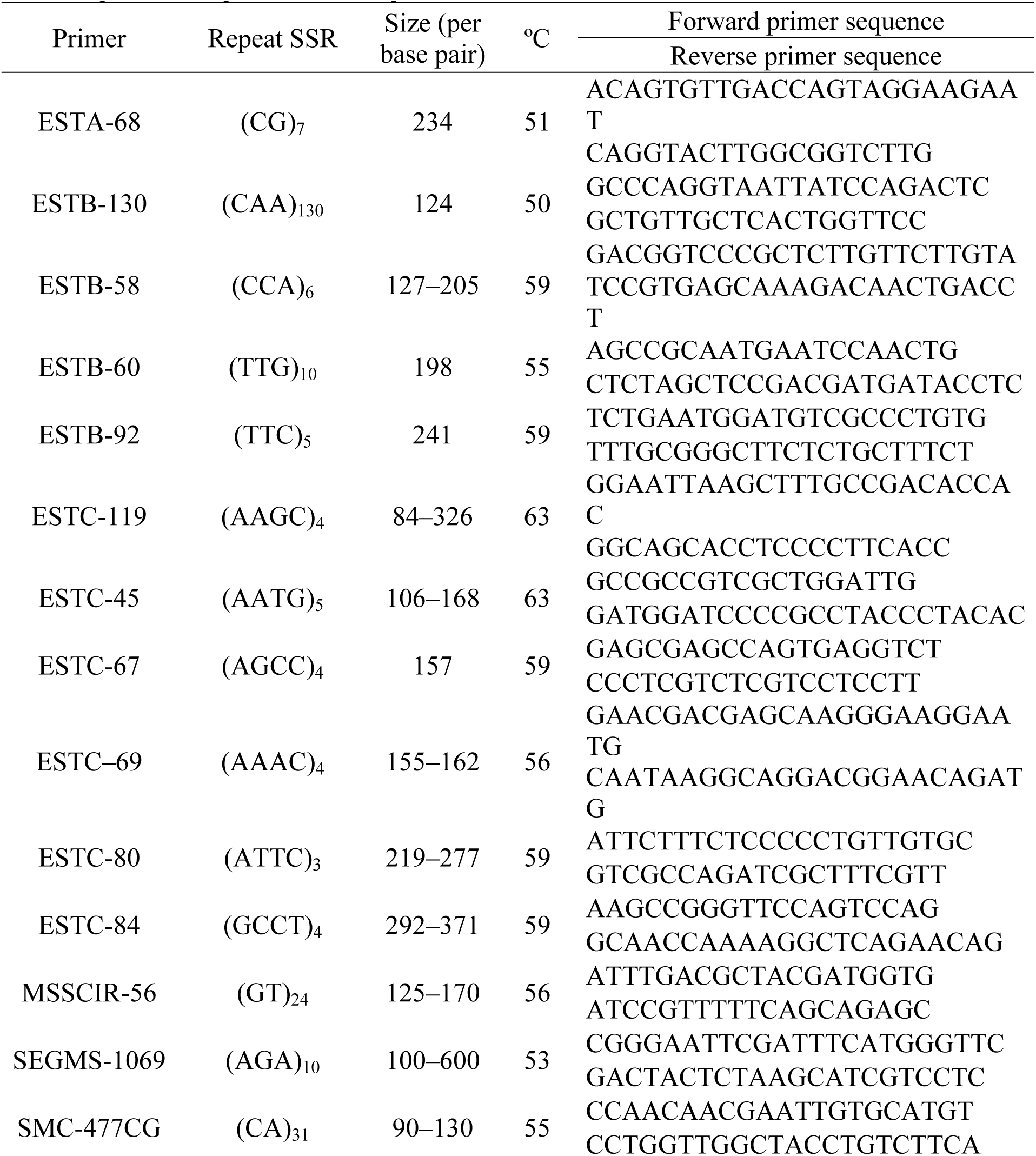
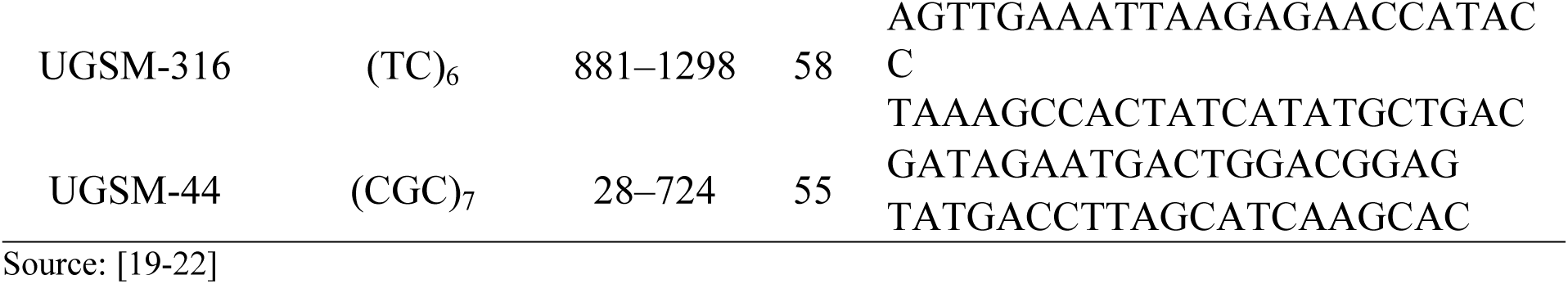
Primers used for genomic DNA amplification of the 12 sugarcane families and their respective amplification temperatures.

TD-PCR offers a high amplification yield when using SSR primers. For example, those with high guanine and cytosine contents (G + C = 60%) present amplification challenges, and the TD-PCR program helps overcome this issue without the need for primer redesign or lengthy optimization [23,24]. Standardization of the PCR reaction is crucial to avoid incorrect products, thereby compromising the accurate characterization of *Saccharum* spp. genotypes.

The observed genetic similarities among the 12 sugarcane families evaluated in this study were significant. The AMOVA indicated that the percentage of genetic variation among families was 28% and that within families was 72% (Table 2). This demonstrates that the high genetic variation detected within families can be utilized in breeding programs and is an important factor in guiding the selection processes for morpho-agronomic traits of interest in crops. Understanding the distribution of genetic variations among and within populations is essential for the effective application of selection methods aimed at genetic enhancement.

**Table 2.** Analysis of molecular variance considering 12 families in the analysis among 120 sugarcane individuals.

| Source | MS | % | LSP |
| --- | --- | --- | --- |
| Among families | 0.5501 | 28% | 0.001 |
| Within families | 0.1155 | 72% |  |
| Total |  | 100% |  |

MS: mean square; LSP: least significant probability.

Forty-two alleles were obtained for the 16 primer pairs (5 SSRs and 11 EST-SSRs) used to amplify the DNA of plants from the 12 sugarcane families. The PIC values for all EST-SSR and SSR markers ranged from 0.25 (UGSM-44) to 0.67 (ESTB-130), with an average of 0.53, indicating that most markers had high discriminatory power and were useful for the study of genetic diversity. The results of our study were in accordance with the results reported by [25], who found an average PIC of 0.67 for 25 EST-SSR and 25 SSR primer pairs in 59 sugarcane genotypes. Similarly, [26] reported a mean PIC value of 0.55 for 158 microsatellite markers in a study.

## Discussion

As previously mentioned, although the majority of sugarcane genetic diversity studies focus on characterizing established commercial varieties or elite individuals, both of which originate from clones, this study examined variability directly from caryopses. Evaluating genetic diversity at this initial stage is crucial to capture the true baseline recombination potential of the population before any directional clonal selection or vegetative propagation occurs.

The families exhibited high degrees of similarity. The lowest similarity was recorded between Families K and D, with a value of 0.8722, while the highest similarity was observed between Families E and I, with a value of 0.9878. In sugarcane, most genotypes used in breeding programs are hybrids resulting from the selection of the best individuals, and the same variety is often used to form several new hybrids. High similarities may result from crosses between closely related ancestors with small genetic distances between them. In this context, it is essential for breeders to understand the genetic divergence among these families and to be cognizant of the genetic resources available for the development of new crosses, particularly for investigating heterosis in new hybrids.

Based on the morpho-agronomic characteristics presented in Table 3, the K family was distinct from the other groups. This differentiation was marked by a notably high ton of stalks per hectare (TSH) and a significantly greater number of stalks per meter, coupled with a considerably lower °Brix compared to that of the other families, thereby justifying the formation of an isolated group. The second subgroup, comprising Families G, J, and L, exhibited very similar averages, particularly in terms of stems per meter (NPM) and average stem diameter (ASD), with nearly identical values. Consequently, the grouping was considered satisfactory from a phenotypic standpoint. Some numerical discrepancies may result from environmental variations that directly influence the specific characteristics.

**Table 3.** Phenotypic evaluation of characteristics stems per meter (NPM), average stems diameter (ASD), °Brix, and ton of stalks per hectare (TSH).

| Families | NPM | (ASD) | °Brix | TSH |
| --- | --- | --- | --- | --- |
| A | 7 | 24.60 | 22.3 | 64 |
| B | 7 | 27.05 | 22.9 | 75 |
| C | 6 | 29.15 | 20.2 | 78 |
| D | 6 | 29.04 | 20.5 | 74 |
| E | 6 | 27.00 | 24.2 | 50 |
| F | 9 | 24.02 | 23.9 | 88 |
| G | 7 | 27.80 | 20.5 | 77 |
| H | 6 | 23.16 | 14.2 | 61 |
| I | 9 | 27.78 | 16.3 | 71 |
| J | 8 | 25.18 | 18.5 | 64 |
| K | 12 | 24.97 | 13.5 | 91 |
| L | 8 | 23.62 | 19.4 | 61 |

Genetic diversity studies in sugarcane conducted by several authors [27–31] treated SSR and EST-SSR markers as dominant, assuming the data as binary (1 for presence and 0 for absence of bands), and they used like those used by [32] and [33], which were developed for diploid organisms, for analysis. The complex genetic structure of sugarcane, which is marked by a high level of ploidy, renders microsatellite markers less effective for these coefficients.

Microsatellite markers have a codominant inheritance with a very informative nature, and analyzing data as binary may lead to the loss of information due to incorrect data analysis, as treating them as binary loses the ability to combine SSR markers, and they are treated as dominant. Specific distance measures that consider ploidy levels are required to properly use microsatellite markers as codominant markers in polyploid species.

According to [34], advanced methods in breeding programs involve the use of molecular markers in diploid species. However, for individuals with different ploidy levels, the use of molecular markers requires analyses that are different from those used in diploid species, such as the calculation of genetic distance, as suggested by [15]. The genetic distance suggested by [15], obtained from microsatellites, considers mutation processes, which enables intrafamily analysis even in polyploid species. In contrast, the Bruvo distance was not used to assess the differences between families. In this study, the RST distance proposed by [35] was used. The distance for microsatellite markers was conceptualized based on the assumption of a stepwise mutation model (SMM), which facilitates the analysis of polyploid species. Thus, these distances are more accurate for studying genetic diversity in highly polyploid species with different ploidy levels, and they enable the interpretation of microsatellite markers as codominant in nature, not as dominant, thus leading to information gain per locus.

Clustering based on the RST coefficients generated a dendrogram (Fig 1), which was validated by the cophenetic correlation test with a value of 0.88 and a significant difference using the Mantel test for the 12 sugarcane families, demonstrating a satisfactory fit between the dendrogram and the similarity matrix.

**Fig 1.**
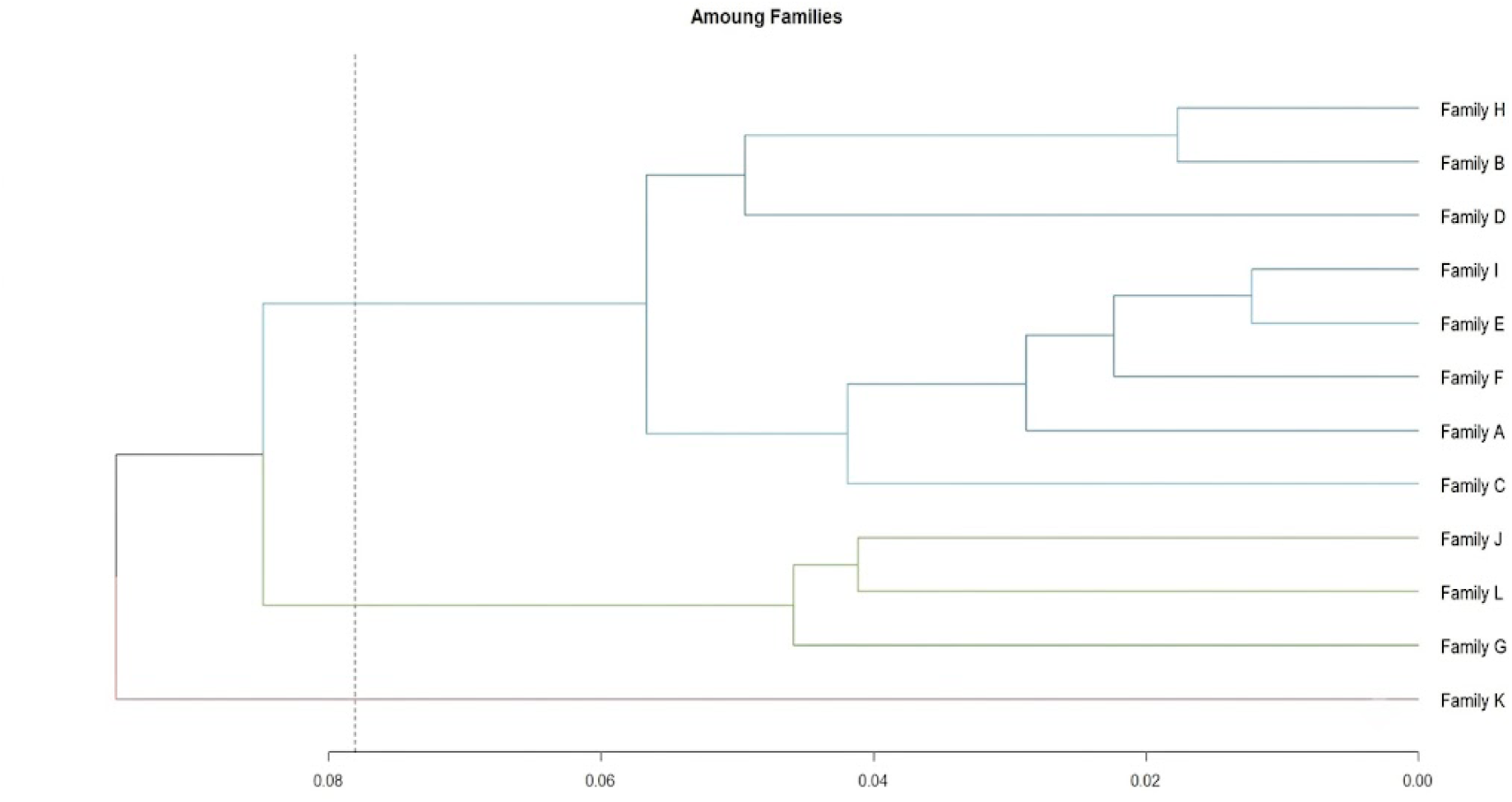
Dendrogram of the 12 sugarcane families. Hierarchical clustering based on RST genetic distance coefficients. The vertical dashed line represents Mojena’s cut-off threshold (0.078), dividing the families into three distinct color-coded clusters: Group 1 (red branch: Family K), Group 2 (green branches: Families G, L, and J), and Group 3 (blue branches: Families C, A, F, E, I, D, B, and H).

Mojena’s cut at 0.078 resulted in the formation of three distinct groups. Group 1 comprised Family K, represented in red. Group 2 consists of Families G, L, and J, which are depicted in green. Group 3 consisted of Families C, A, F, E, I, D, B, and H, as shown in blue. The isolated group formed by Family K illustrated the genetic distance of this family and opened the perspective that crossing Family K with others could broaden the genetic diversity of other genotypes used in breeding programs. [12] pointed out the high levels of genetic similarity among sugarcane varieties, leading to the conclusion of some authors [36–38] that sugarcane has a narrow genetic base. [9] reported that a narrow genetic base is a characteristic of modern sugarcane varieties because of the limited number of parents involved in sugarcane breeding. The limited genetic diversity of contemporary sugarcane cultivars can be attributed to their derivation from fewer than 20 genotypes originating from breeding programs in Java and India [39]. However, [4] ascribed the narrow genetic base to successive backcrosses to insert characteristics such as disease resistance.

The 120 sugarcane individuals analyzed with the 16 microsatellite loci using Bruvo’s distance based on allele frequencies were compared, and the dendrogram constructed using the WPGMA clustering method indicated high genetic variability within the families (Fig 2).

**Fig 2.**
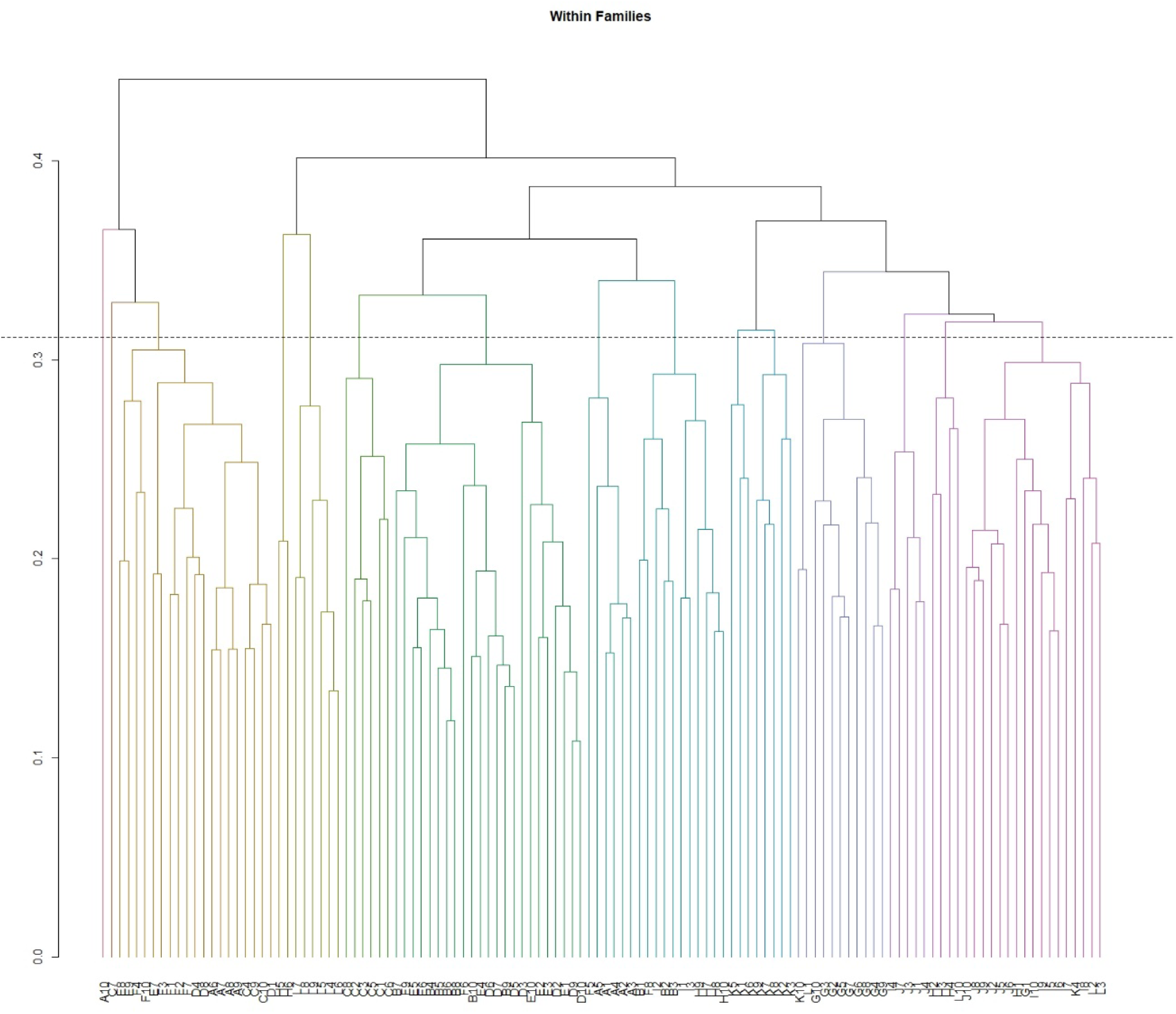
Dendrogram of the 120 individuals from the 12 sugarcane families. Hierarchical clustering generated by the WPGMA method based on Bruvo’s genetic distance using 16 microsatellite loci. The horizontal dashed line indicates Mojena’s statistical cutoff threshold (0.3110), which differentiates the 15 color-coded groups and highlights the high intrafamily genetic variability.

When the Mojena statistical cutoff (0.3110) was used, 15 groups were formed, indicating that the degree of similarity varied from 0.3735 (L5 × C2) to 0.8915 (D9 × D10).

Currently, hybrid sugarcane cultivars are predominant, and genetic variability is obtained from crosses of divergent individuals, consequently reducing genetic narrowing and vulnerability [40,41]. Low degrees of similarity were observed between L5 × C2 (0.3110), F4 × J4 (0.3857), and F4 × J6 (0.4123). These crosses can be maximized in breeding programs, as they tend to generate descendants with low similarity, increasing their variability in the next stages of breeding. This is because the chance of selecting individuals with desirable agronomic characteristics, high heterosis, and low similarity is greater than that of crossing individuals with high similarity.

Although sugarcane is cited as a species with a narrow genetic base, the results of the present study, which analyzed five SSR and 11 EST-SSR loci in 12 families from the RIDESA breeding program, indicated high genetic variability within these families. The sugarcane genome is notable for its extensive polyploidy, containing approximately 8–14 copies of homologous chromosomes and displaying highly heterozygous loci. Genomic organization has significant potential to generate diverse genetic combinations, potentially leading to population segregation [42]. This considerable variability challenges the assumption that sugarcane has a limited genetic basis. The genetic basis can be regarded as broad, but remains underexplored. The generation of highly contrasting individuals within the same family indicates significant variability within the *Saccharum* spp. genome, suggesting that further exploration is needed.

In the present study, the sugarcane population was formed by 12 families of hybrids, which were analyzed in the first generation; that is, they were sexually propagated via seeds, which may explain the large genetic variation within these families. This higher percentage of genetic variation detected within the families, with 72% of the variation within, reflects a higher variability in the population, enabling crossbreeding between contrasting individuals with low similarity to generate variability, as may be the case for the cross between individuals L5 × C2 (0.3110) and J4 × F4 (0.3857), to generate new heterotic groups. Significant variation within families permits a reduction in the number of families used. As a result, increasing the number of seeds per family may be required to enhance genetic variability.

Clustering analysis employing a model-based Bayesian algorithm demonstrated that genetic variability was lower among the 12 families than within the families. Twelve crosses between 19 different genotypes generated only two subpopulations, indicating that the divergence between the 12 families was low. The optimal K value determined using Bayesian analysis grouped the 120 sugarcane plants into two subpopulations (DK1 = 0.00, DK2 = 438.1319, DK3 = 0.64451, DK4 = 48.7293, and DK5 = 0.09223). The bar plot obtained for the K value (K = 2; DK = 438.1319) was consistent with the evidence that the 120 plants from the 12 families formed two genetically structured populations. In the clustering, the colors represent two different clusters corresponding to the differential proportions of plants in each group (Fig 3). The bar plot indicates that 90% of the sugarcane plants in the red group exhibited allele frequencies, with 11% of these plants sharing frequencies with those in the green group. Conversely, 89% of the sugarcane plants in the green group displayed allele frequencies, with 10% sharing frequency with the plants in the red group. Each bar in the graph represents a plant and its inferred proportion of genomic admixture. Individual plants sharing allele frequencies in the red and green groups revealed a genome admixture. The genome admixture may be due to the common genotypes used in the crosses to obtain the 12 families and/or from the common parents of some of the genotypes used in the crosses.

**Fig 3.**
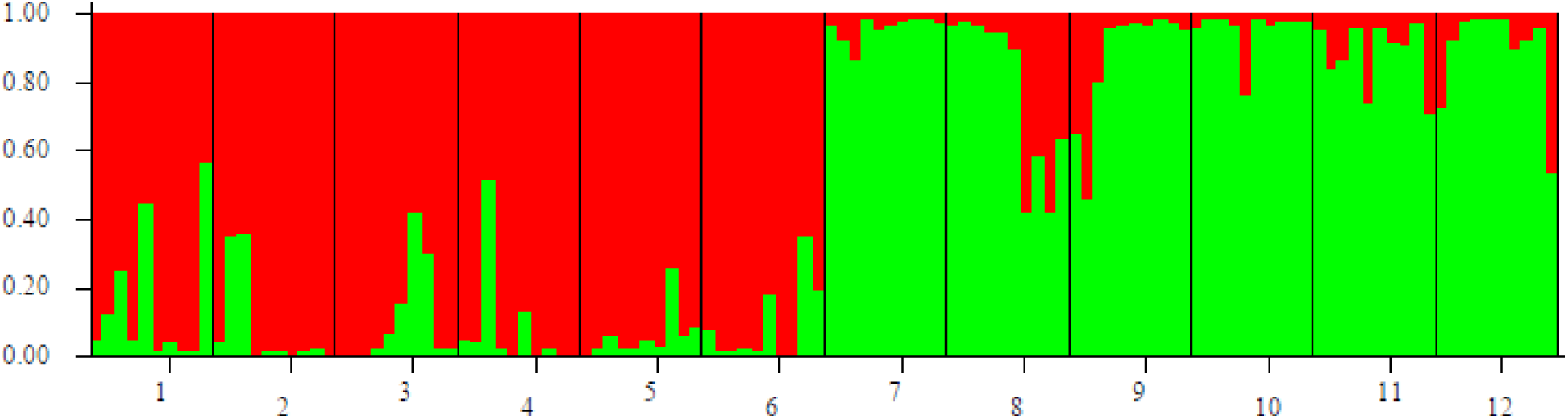
Population structure and genome admixture of sugarcane families. Bar plot generated from genetic clustering analysis (K=2). Each vertical bar represents an individual plant, indicating its inferred proportion of genome ancestry assigned to the red or green genetic clusters. Numbers 1 to 12 along the x-axis correspond to the 12 evaluated families, separated by vertical black lines.

The results obtained in the present study using 16 pairs of microsatellite primers showed a wide genetic variety to be explored within the 12 families of sugarcane analyzed. Microsatellite markers have shown high efficiency in assessing genetic variability in *Saccharum* spp. and discriminating genotypes being an excellent tool for plant breeding.

## Materials and Methods

The experiment was conducted at the Irrigation Technical Center (CTI), an entity affiliated with the Department of Agronomy (DAG) at the State University of Maringá (UEM), located in Maringá City, Paraná (coordinates 23°25’57” S, 51°57’08” W, and 542 masl). This area features soil classified as Dystroferric Red Nitosol with a moderate A horizon and clayey texture [43]. The experimental units consisted of ten sugarcane seedlings in each planting furrow, originating from the caryopsis. These units were arranged in an augmented block design [44] with an effective area of 7 m² per furrow, with no repetition and two commercial checks, RB986928, and RB867515. This design was utilized because sugarcane genotypes originate from caryopses, leading to each plant being unique, even though they belong to the same family. The plant material comprised 120 individuals from 12 full-sib sugarcane families sourced from the RIDESA Genetic Improvement Program (Table 4). These individuals were selected based on the agronomic traits listed in Table 3.

**Table 4.** Selected crossings for genetic divergence studies in the 12 families of sugarcane.

| Family | Crossings (♀ x ♂) |
| --- | --- |
| A | CTC9 × RB946903 |
| B | RB036088 × TUC74-15 |
| C | RB036152 × RB036088 |
| D | RB106802 × RB855036 |
| E | RB986952 × RB986960 |
| F | TUC71-7 × RB036152 |
| G | CTC-15 × RB036145 |
| H | CTC-9 × RB965518 |
| I | RB036122 × RB03611 |
| J | RB946015 × RB936109 |
| K | RB97319 × RB966928 |
| L | RB986960 × RB966952 |

Young leaves were collected from each plant originating from caryopsis, placed in aluminum packages, frozen in liquid nitrogen for five minutes, and stored in an ultra-freezer (−80 °C) until used for DNA extraction. Genomic DNA was extracted using the protocol described by [45], with some adaptations for sugarcane. For DNA amplification, primer pairs were selected from a set of 49 microsatellite primers that make up the primer bank for sugarcane, according to [19–22]. At the end of the tests, 16 primer pairs that successfully amplified the DNA and presented satisfactorily defined DNA fragments on the electrophoresis gel were selected (Table 1) and used for the analysis of the 12 *Saccharum* spp. families (Table 4).

DNA samples from sugarcane families were amplified using a VeritiTM DX 96-Well Thermal Cycler (Applied Biosystems). Touchdown-PCR (TD-PCR) software was employed to amplify DNA using 15 of the 16 primer pairs, as detailed in [46]. For a specific primer pair, an annealing temperature of 53 °C was employed.

The amplified DNA was evaluated using a 4% agarose gel, composed of 50% ultrapure agarose and 50% MetaphorTM agarose, and 0.5X TBE buffer (89 mmol·L^-1^ Tris, 89 mmol·L^-1^ boric acid, and 2 mmol·L^-1^ EDTA) for separation of the amplified DNA segments, applying 60 Volts for 4 h. A 100 bp ladder (Invitrogen) was used as a molecular-weight marker. Following electrophoresis, the gel was stained with 0.5 µg·mL^-1^ ethidium bromide for 1 h and then photographed using a UV transilluminator (Loccus Biotecnologia) and the L-Piximage software.

The DNA segments amplified using PCR were identified, and the base-pair molecular weights were calculated to generate a matrix for each gel. Gel readings were performed using the *ImageJ* software (Java 8 [64-bit], National Institutes of Health, Bethesda, USA).

After constructing a matrix with the number of base pairs for each fragment, *R Software* [47] was used for all analyses. The genetic parameters analyzed included allele frequencies per allele per locus, the mean of the highest allele frequencies, and the polymorphism information content (PIC) proposed by [48] using equation 1:

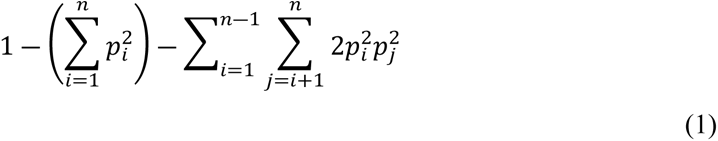

where p_i_ and p_j_ are the allele frequencies for alleles i and j, respectively, and n is the total number of alleles.

The genetic similarity between individuals was calculated using Bruvo’s genetic distance [15]. The RST distance [35] was used to calculate the distance between the families. The distances were computed from allele frequencies estimated using the method proposed by [49], with the bootstrap resampling technique with 5,000 permutations. All analyses were conducted using the Polysat package version 1.7-7 [50].

A dendrogram was constructed using the weighted pair-group method using the arithmetic averages (WPGMA) hierarchical clustering method. Cophenetic correlation [51] was used to validate groupings. Subsequently, the significance of the groupings was tested using the Mantel test [52] with 10,000 permutations. Group separation was established using Mojena’s statistical criterion [53] based on the relative size of the fusion levels (distances). Analysis of Molecular Variance (AMOVA) [54] was performed using the ADEGENET package version 2.1.10 [55], with 50,000 iterations.

The genetic structures of the 12 families were evaluated using STRUCTURE version 2.3.4 [56]. The genotypes were clustered, with the number of clusters (K) ranging from 1 to 12, and tested using the admixture model with a burn-in period of 50.000 repeats followed by 100.000 Markov Chain Monte Carlo (MCMC) repeats, considering the presence and absence of alleles across the sample. The true number of populations (K = 2) was often identified using the maximal value of Δ (K) returned by the software (Fig. 4).

**Fig 4.**
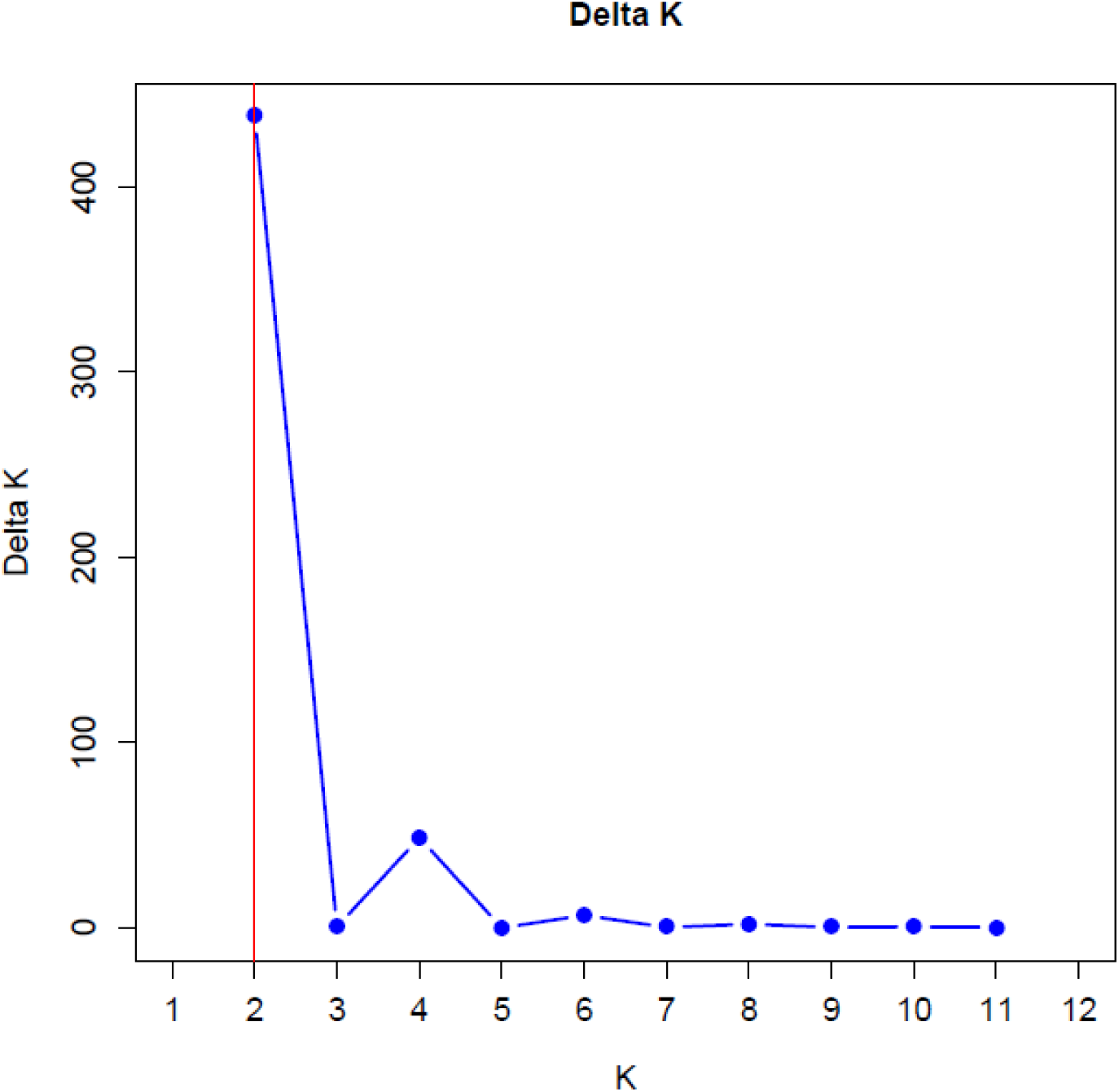
Delta K plot for determining the optimal number of genetic clusters (K). Estimation of the true number of sub-populations using Evanno’s method. The highest peak along the blue line, highlighted by the vertical red line, identifies K=2 as the optimal number of genetic clusters for the analyzed sugarcane families.

The most probable number (K) of subpopulations was identified as described by [57]. The graphical output display of the STRUCTURE results was used as input data using StructureSelector, a web-based software used to visualize the STRUCTURE output and implement the Evanno method [58] to display a graphical representation.

Despite the high genetic similarity among the 12 sugarcane families in the present study, the high genetic variation detected within these families can be used in breeding programs to generate new crosses. The substantial genetic variability within families can effectively guide the crossing of specimens from different families. This method aims to regulate genetic variability throughout the selection process for morpho-agronomic traits and disease resistance characteristics of interest in the culture.

Furthermore, our findings reveal that the genetic base of sugarcane is broad, even among individuals with common ancestors, enabling crosses with low similarity that actively enhance population diversity. Consequently, to maximize genetic gains, sugarcane breeding programs should optimize their resources by increasing the number of individuals evaluated per family while reducing the total number of families, as the primary source of genetic variability resides within the families rather than between them.

## Acknowledgements

The contribution of materials for this study by the Interuniversity Network for the Development of the Sugarcane-Energy Sector (RIDESA) is duly acknowledged.

